# Transmission analysis of a large TB outbreak in London: mathematical modelling study using genomic data

**DOI:** 10.1101/761411

**Authors:** Yuanwei Xu, Hollie Topliffe, James Stimson, Helen R. Stagg, Ibrahim Abubakar, Caroline Colijn

## Abstract

Outbreaks of tuberculosis- such as the large isoniazid-resistant outbreak centered on London, United Kingdom, which originated in 1995- provide excellent opportunities to model transmission of this devastating disease. Transmission chains for tuberculosis are notoriously difficult to ascertain, but mathematical modelling approaches, combined with whole-genome sequencing (WGS) data, have strong potential to contribute to transmission analyses. Using such data, we aimed to reconstruct transmission histories for the outbreak using a Bayesian approach, and to use machine learning techniques with patient-level data to identify the key covariates associated with transmission. By using our transmission reconstruction method that accounts for phylogenetic uncertainty, we are able to identify 24 transmission events with reasonable confidence, 11 of which have zero single nucleotide polymorphism (SNP) distance, and as maximum distance of 3. Patient age, alcohol abuse and history of homelessness were found to be the most important predictors of being credible tuberculosis transmitters.

## 1. Introduction

Analyses of chains of transmission- i.e. who infected whom- are critical tools within outbreak control. In tuberculosis (TB), transmission analysis is particularly challenging because TB has the potential for dormancy in infected individuals many years after transmission, making it hard to distinguish recent transmission from reactivation. Additionally, in low-incidence, wealthier, nations, the disease is often concentrated in populations that are under-served by traditional healthcare models, resulting in infectious individuals taking many months to be diagnosed. Within the public health process for patients with infectious respiratory disease, it can be challenging to identify an individual’s contacts over long periods of time, particularly within under-served population groups, who may have unstable housing and mistrust traditional systems of authority.

Since the advent of next-generation sequencing technologies has made this feasible [20], whole genome sequencing (WGS) data are increasingly gathered in efforts towards TB control, and there have been high hopes that WGS technologies will greatly facilitate outbreak reconstruction. In high-income countries WGS data and demographic and epidemiological data are now often gathered for TB. National TB control programs following WHO guidelines collect a standard set of data including demographic and clinical data along with data on treatment outcomes and bacteriology [19, 14], some of which are likely to be related to transmission. Local programs may collect further variables, which can be crucial in controlling and eliminating outbreaks [19]. In 2017, England was the first country worldwide to roll out routine WGS for TB cases [17]. It is not clear to what extent WGS data will reveal transmission events, though it is now established that sequences alone, at least with current bioinformatics analysis pipelines, are insufficient to reliably determine precisely who infected whom [5, 6, 7, 4]. But the context of increased availability of WGS data, together with demographic and clinical covariates, provides researchers with new challenges – to what extent can incorporating demographic and clinical data with WGS aid in understanding transmission?

Within London, particularly the north of the city, a long-standing outbreak of isoniazid (INH) mono-resistant tuberculosis, first defined by a shared RFLP cluster and later defined by a shared identical 24-loci MIRU-VNTR type, has existed since 1995 [16, 13, 11, 5]. By 2013, there were 501 cases in total in the UK. Extensive contact tracing and transmission analysis were done in the first years following detection of the outbreak in 2000 [16, 13]. The outbreak has been of particular concern; there have been hundreds of cases and it has contributed to rising isoniazid resistance in England [13]. The outbreak has showed signs of high transmissibility- with only brief contact sufficient for transmission [16, 5]- and a high proportion of smear-positive cases [11, 5]. During and after the outbreak, retrospective outbreak questionnaires and patient interviews were completed by TB clinic nurses, gathering information such as drug and alcohol use, history of homelessness or imprisonment and treatment history. Recently, isolates from the outbreak cluster were sequenced with whole genome sequencing (WGS) to aid in resolving the transmission network, but due to low levels of detectable variation, individual transmission events could not be reliably inferred [5].

Here, we combine WGS data, data on times of sampling, demographic, clinical and other host data to analyse this complex outbreak. We first construct timed phylogenetic trees using WGS data together with sampling times. We introduce a new approach to reconstructing individual transmission events, jointly analysing a posterior collection of timed phylogenetic trees while sharing key model parameters. This takes phylogenetic uncertainty into account while constraining reconstructed transmission events on different posterior phylogenies to have the same underlying epidemiological parameters. This analysis allows us to estimate how many unsampled cases there were, how long individuals took from original infection to infecting others, and the time between initial infection and sampling, taking phylogenetic and parameter uncertainty into account. Finally, we relate the extended demographic and clinical data to transmission by training machine learning tools to predict which individuals were likely transmitters, using the covariate data alone.

## 2 Methods

### 2.1 Data

The London isoniazid resistant TB outbreak is characterised by Public Health England as cluster E1244 (strain type 424332431515321236423–52 and including an untypeable 3690 locus). Previous work documents the data collection, surveys and questionnaires, contact tracing and WGS used as part of outbreak investigations [16, 13, 11, 5]. The cluster was first identified using IS6110 RFLP analysis, by screening method based on PCR, and then using a unique 24-locus MIRU-VNTR type [5]. In [11] cases were defined as part of the outbreak if they had an INH-monoresistant strain, were diagnosed between 1995 and 2006, had the RFLP or MIRU-VNTR pattern matching the outbreak, and were either a resident of London or had epidemiological links with London [11]. The outbreak then continued after 2006 and was described with sequencing data by Casali et al [5].

Covariates including variables on homelessness, prison links, drug and alcohol usage were obtained from the patient surveys, questionnaires and interviews along with medical records; we visualise some of these data in Figure 1c. We take the following approach to missing data: for categorical variables with two strata (e.g. “yes” and “no”; this describes most of our variables), if a varaible was missing more than 40% of its data, then the missing values were replaced by “Unknown”. For all other variables, the R package mice was used for multivariate imputation. In doing so, we had assumed that the missing data was missing at random. The decision to replace rather than impute the missing data is based on the observation that if many entries are missing, there may not be enough information for imputation, and so the result could be far from the truth. On the other hand, discarding the variable completely will result in a loss of information, and we wish to use the data that are available.

**Figure 1:**
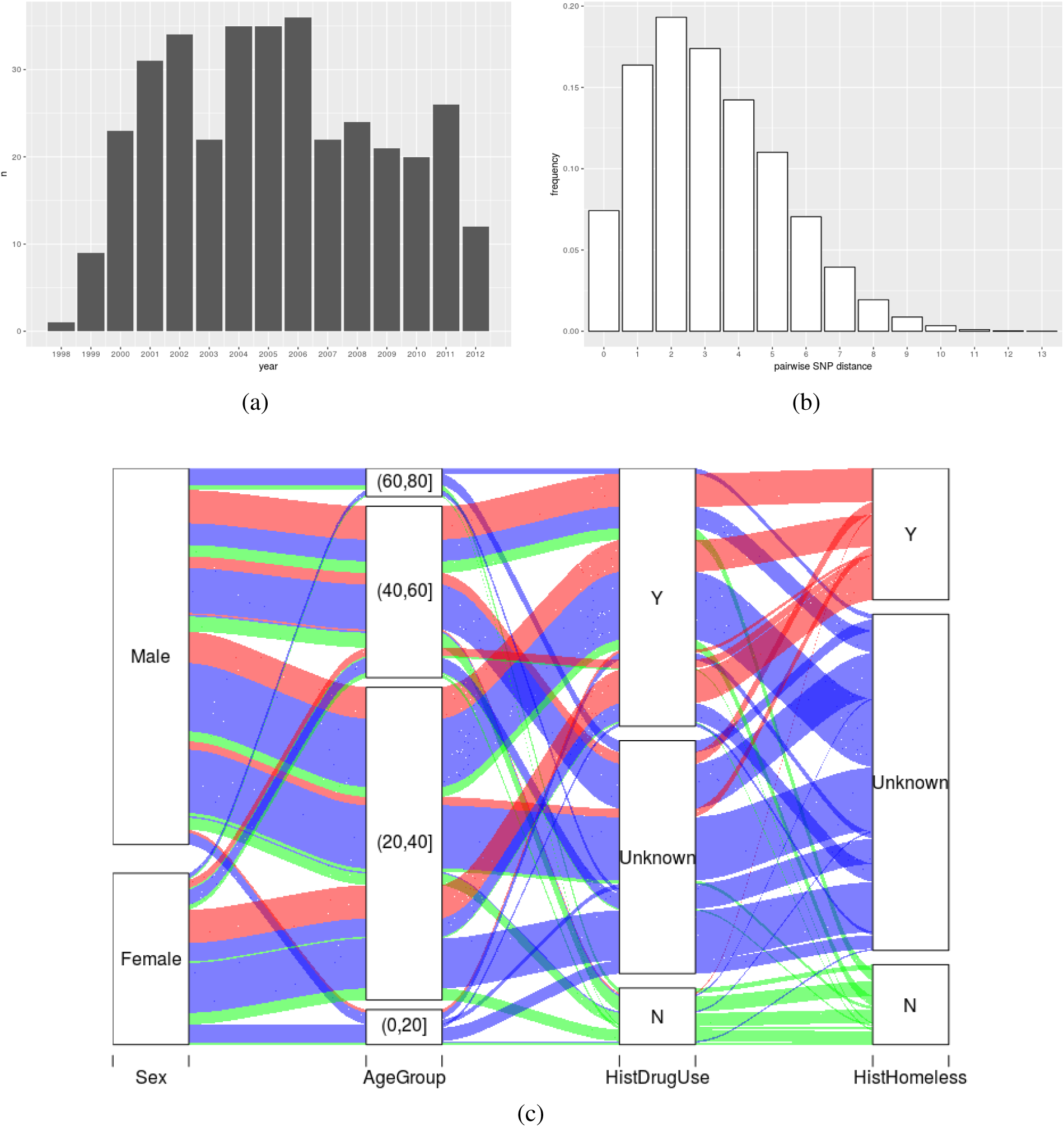
(a): Number of sequences in our dataset by year. (b): Frequency of pairwise SNP distance. (c) Illustration of some of the covariates in an alluvial plot showing how many individuals are in each category and how many share categories from one column to another. Colours correspond to homelessness history: Y—red; N—green; Unknown—blue. As an example of the interpretation of the plot, nearly all those who have a ‘yes’ for a history of homelessness (red) either also have ‘yes’ or ‘unknown’ for a history of drug use (red bands reaching from Y and Unknown in the HistDrugUse column up to the Y category for HistHomeless).

### 2.2 SNP calling and phylogenetic reconstruction

Isolates were cultured and then whole genome sequenced [5] on an Illumina HiSeq with a read length of 100bp at the Wellcome Trust Sanger Institute; raw data are available in the European Nucleotide Archive under the accession number ERP003508. Samples in this study were excluded from the analysis if any issues were recorded with their culturing in the lab, such as lack of growth, contamination or other reasons potentially impacting quality. An assessment of sequence quality is initially carried out using FastQC. Raw fastq reads were then filtered for length and trimmed for low-quality trailing base pairs using Trim Galore; any trimmed reads which were shorter than 70 base pairs were discarded. Reads were aligned to the H37Rv NC000962.3 reference genome using the BWA MEM algorithm, with duplicate reads removed using Picard’s MarkDuplicates tool. SNPs in hypervariable regions, repeat regions, and mobile elements were excluded. Local realignment round insertions and deletions (indels) were carried out using the GATK IndelRealigner tool. SNPs were identified using FreeBayes with a minimum mapping quality of 30 and minimum base quality score of 20. Isolates with a high proportion of apparent mixed or heterozygous SNP calls (i.e. those with more than 20% reads supporting the reference allele) were excluded from analysis. A variable-site alignment was created as a fasta-format multiple sequence alignment which excludes non-variant bases, along with an index mapping the base number of the alignment to the corresponding location on the reference genome. Calls made with a read depth of less than 30 across all the samples in the study were also excluded.

The phylogenetic tree-building software BEAST [3] was used to build timed phylogenetic trees, using a constant population coalescence model, generalised time reversible substitution model and strict molecular clock. Instead of trying to obtain a single optimal timed phylogenetic tree from the posterior, we sample a collection of them. This ensures that we capture as much diversity as possible from the BEAST posterior, to achieve robust uncertainty quantifications in our subsequent analysis.

### 2.3 Transmission inference

We perform Bayesian inference of transmission trees given a timed tree using the TransPhylo approach [7], but we extend the approach to simultaneously infer transmissions on a subsample of BEAST trees rather than using just one. We base all downstream analysis on a combined set of posterior transmission trees inferred from these distinct phylogenetic trees. We also allow the flexibility of sharing model parameters across different input phylogenetic trees, so that only a single parameter set is updated instead of *N* sets for *N* timed phylogenetic trees. This results in better mixing for the underlying Markov chain Monte Carlo algorithm than not sharing parameters.

Since TransPhylo assumes that sequences are from unique hosts and it is not designed to handle multiple sequences from the same host, in order to avoid confusion of a host infecting itself, we only keep the earliest sequence from each host in the input phylogenetic trees.TransPhylo was run on the joint tree space of 50 trees from the BEAST posterior, with parameter sharing, for 4 *×* 104 iterations, and output transmission trees were collected every 50 iterations. The generation time and time-to-sampling are both described by Gamma densities with fixed parameters, for generation time, the shape and scale parameters used are 1.3 (unitless) and 2.5 years, respectively; and for the time between infection and sampling, the shape and scale parameters used are 1.1 (unitless) and 6.0 years. The second parameter of the negative binomial offspring distribution was fixed to be 0.5 and not updated. We fixed the within-host coalescent time unit to be 100/365 [8]. A prior belief that case-finding was effective during this outbreak was reflected in TransPhylo with an informative Beta prior for the sampling probability *Π*, with the two parameters being 8 and 2.

### 2.4 Patient-level prediction from metadata

The “ground truth” answers to many questions in an outbreak reconstruction – who infected whom and when – are typically not known. We use the posterior transmission trees from TransPhylo as a proxy for this ground truth. For example, suppose that we are interested in predicting whether a host has transmitted TB. We can describe whether an individual is a “credible transmitter” by setting a binary variable to be true if more than half of the posterior transmission trees suggest that the host infects at least one other; while the true transmission events are unknown this allows us to capture variation in the likelihood that an individual transmitted to another during the outbreak. We could also be more stringent by assigning a true label only when over 80% of transmission trees imply that the host infects someone else; in this case, the resulting true positives will more closely reflect the TransPhylo estimates of who is a transmitter, but false negatives will likely increase as well.

Once we have extracted a host-level variable of interest from the posterior transmission trees produced by TransPhylo, we can then train a machine learning algorithm to predict this target variable, using either 1) both the metadata and other predictors extracted from TransPhylo such as the generation time and time-to-sampling for each host, or 2) the metadata alone. Here we choose the latter because we are interested in assessing whether the covariates in the metadata have predictive power for identifying credible transmitters.

We explored two machine learning tasks: predicting whether an individual is (likely) a transmitter of TB, and predicting whether an individual is estimated to have a longer-than-usual generation time. Accordingly, in the first task, the response was chosen to be a binary variable that is true if the posterior probability that the individual in question infects at least one other individual is greater than 0.5, and false otherwise. A random forest classifier was trained with five-fold cross validation, so that each model was used to predict the data that it has not seen during training. In the second task, we create a new binary variable (’long’ generation time or not). The generation time is estimated from the TransPhylo posterior transmission trees by subtracting the mean infection time of a host from the mean first transmission time of that host (Figure 5c). The mean is taken over all posterior transmission trees in which the host ever infects another (regardless of who they infect, and even if they have low posterior probability of infecting anyone). Naturally, generation times are censored by the end time of the data; individuals infected very recently have had less opportunity to infect others, and any secondary cases we do have in the data will have happened rapidly. We consider the generation time to be ‘long’ if it is greater than 2 years, and short otherwise. We train a random forest classifier as in the first task, using the same set of training and validation data. Using 0.5 as a threshold of probability of transmission, TransPhylo predicts that 202 (61%) cases were transmitters and 127 (39%) cases were non-transmitters. 77 (23%) cases had generation times above 2 years and 252 (77%) below 2 years.

## 3. Results

In the final dataset there are 351 sequences, of which 94 were identical, and 269 variable sites. We used 329 of the sequences for the transmission analysis, as we used only the earliest sequence from each host. We show in Figure 1 the total number of sequenced isolates per year between 1998 and 2012. The frequency of all pairwise SNP distances is shown in Figure 1(b). Among other patterns, we note that if a patient has a history of homelessness, then he or she tends to also have used drugs; most patients are age 20 and 50, with more males than females. There are missing data, which is to be expected, as patients may be unwilling to disclose some of this information, and record-keeping over a long period across multiple sites can be prone to error and loss.

In order to have a picture of the overall transmission network, we show in figure 2 the maximum a posteriori (MAP) transmission tree from the combined TransPhylo posterior. One advantage of TransPhylo is its ability to model unsampled cases and estimate their numbers; here we estimate an average of 53 unsampled cases (Figure S2) compared to 329 sampled, reflecting a relatively high case finding rate. Figure S1 shows the MCMC trace plot of the model parameters. The final 30% of the transmission trees (sampled every 50 iterations) corresponding to each of the 50 timed phylogenetic trees were joined together in a combined posterior that is used for downstream analysis. Because they share parameters, the mean of `off.r`, or equivalently the *R*_0_, for any timed tree is 1.08; and the mean sampling probability `pi` is 0.75. Recall that the within-host coalescent time unit (`neg`) was fixed to be 0.27 (100/365).

**Figure 2:**
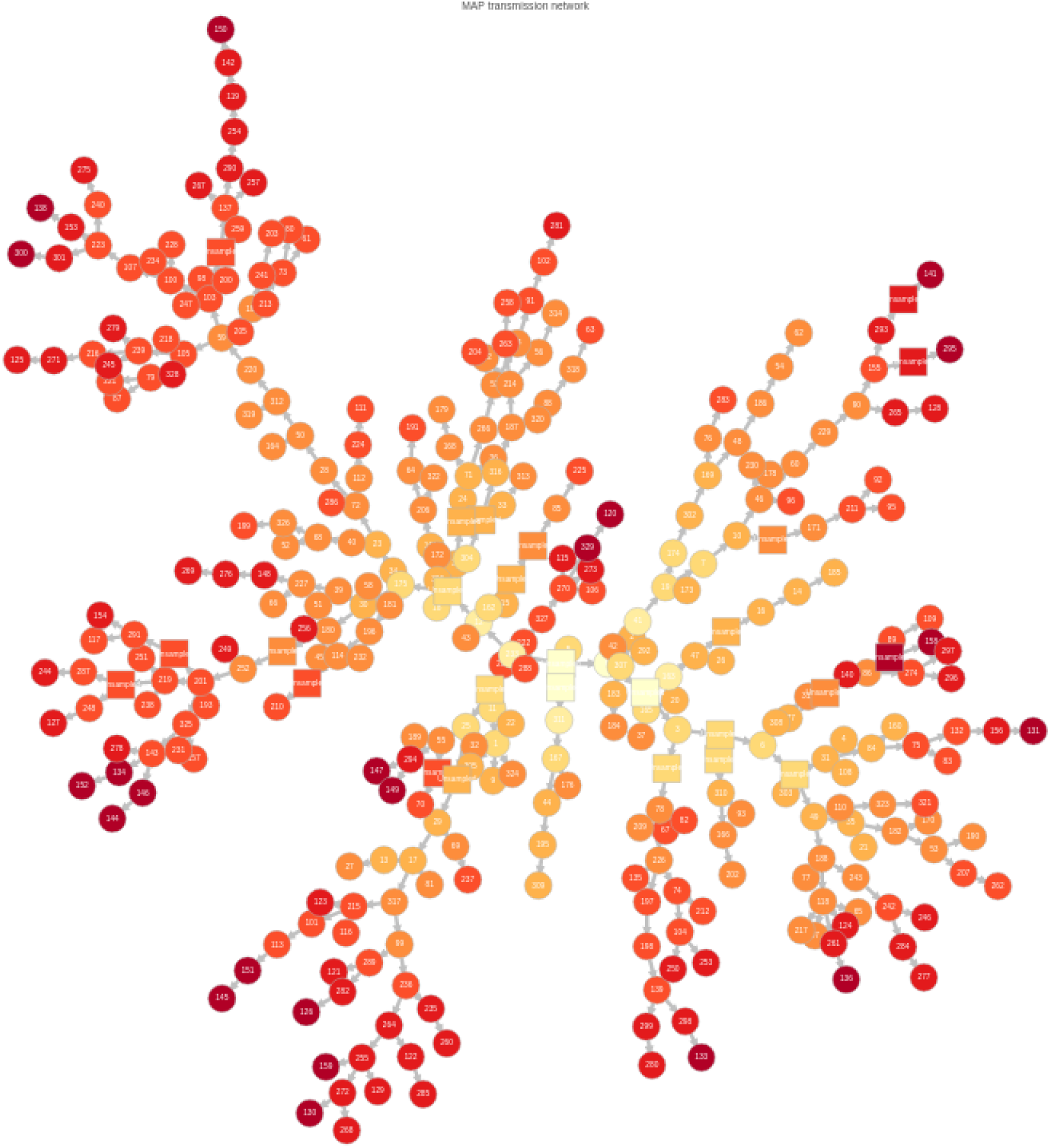
Maximum a posteriori (MAP) transmission tree of the combined TransPhylo posterior. Nodes (hosts) are colored by time of infection and change from yellow to orange to red as infections become more recent. Sampled cases are shown as circles whereas unsampled ones are shown as squares.

In contrast to credible TB infectors (see Methods), we sought credible transmission pairs, namely transmission pairs from individual *i* to *j* that have posterior probability greater than 0.5. There are 16 such transmission pairs: 8 with no SNPs, 5 with 1 SNP, 2 with 2 SNPs, and 1 with 3 SNPs between the host isolates. In addition, we have identified 9 transmission pairs with posterior probability greater than 0.5 in which the infector is unsampled. Previous analysis of this outbreak [5] (Fig. 3) using WGS data identified 5 transmission events (compared to 16 here and an additional 9 with unsampled infectors), though considerable uncertainty remains. There is maximum of 4 SNPs in two of the transmission events suggested by WGS in [5], in contrast to a maximum of 3 SNPs in one of the transmission events identified by our approach. In the 16 pairs we identified, there are 5 pairs where both individuals were treated in the same hospital, 7 pairs where both are of the same ethnicity, 4 pairs where both have been drug users and 5 pairs where both have links with prison. We also note that there is one pair that simultaneously shares all these attributes. Two of our identified pairs are in agreement with known reported contacts, but contact data are not available for the majority of our cases.

**Figure 3:**
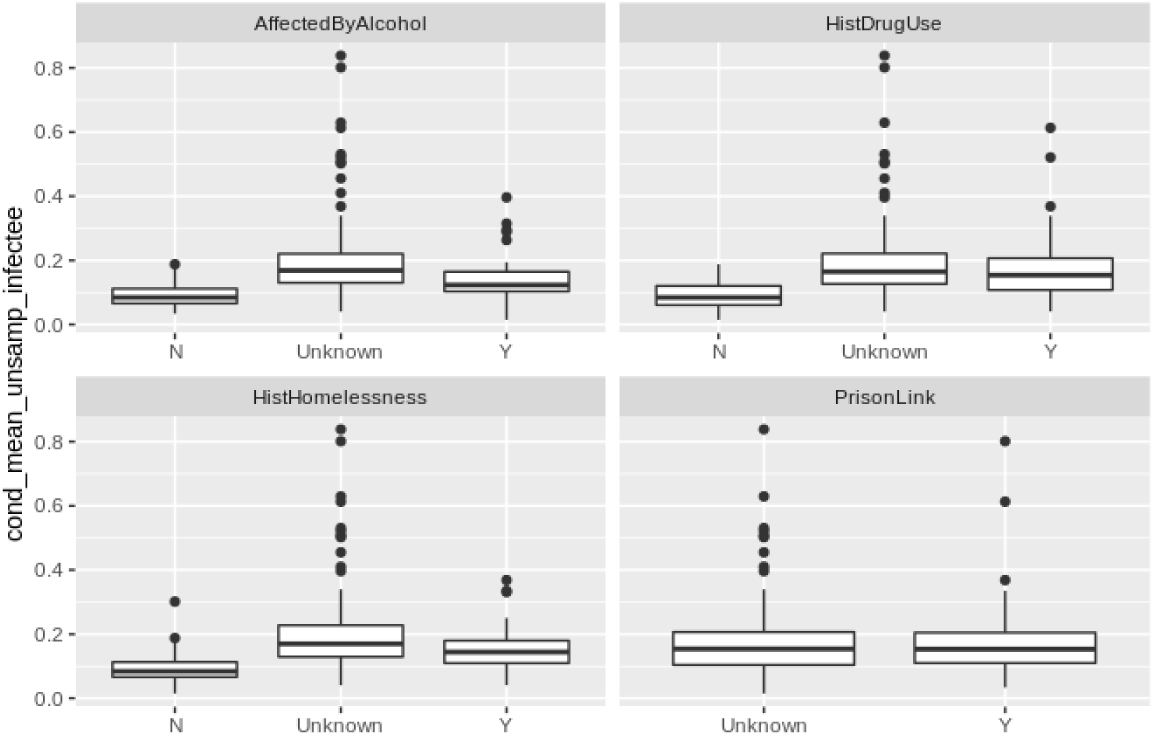
Mean number of unsampled infectees of hosts in different categories defined by four covariates, conditioned on the host infecting at least one other host in the posterior transmission trees.

Even if highly likely pairs are not revealed by WGS data, we can interrogate the Bayesian transmission trees to obtain information that can be useful in outbreak control and case finding. In particular, we explored the relationship between host covariate data and whether hosts are inferred to have infected, or been infected by, unsampled cases.

Because a host can have many infectees but can only have one infector, we compute the mean number of unsampled infectees over the set of posterior transmission trees in which the host infects at least one other host, and the probability of having an unsampled infector, conditioned on the host not being the index case. We group the estimates by four covariates, shown in figure 3 and 4. We find that an individual tends to infect more unsampled cases if they have been affected by alcohol or drugs, or have a history of homelessness. Based on our data, we can not conclude whether having prison link is connected to having more unsampled infectees, because the categories in our prison data are ‘yes’ or ‘NA’ (unknown). The plot of the probability that an individual’s infector is unsampled shows a similar pattern, i.e., an individual is more likely to have an unsampled infector if he or she has used drugs or alcohol, or has been homeless.

**Figure 4:**
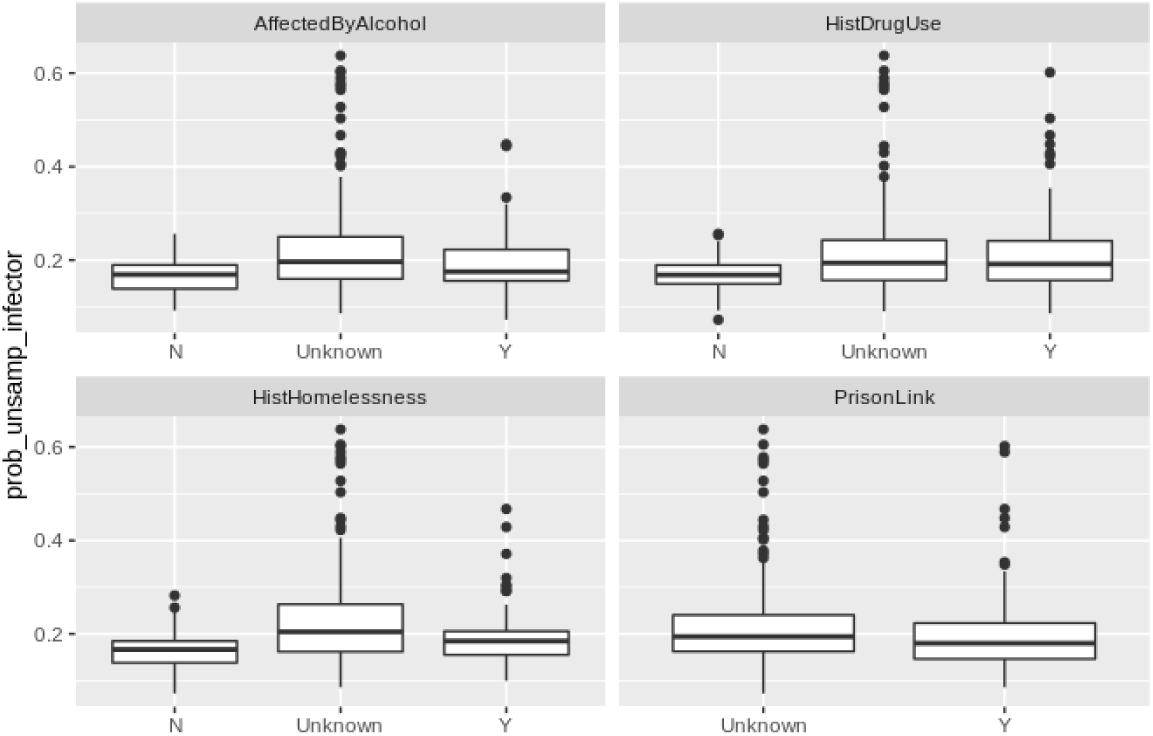
Probability of a host having unsampled infector in different categories defined by four covariates, conditioned on the host not being the index case in the posterior transmission trees.

Our outbreak reconstruction with WGS data can also help interrogate the timing of transmission and sampling in reconstructions consistent with genomic data, despite the fact that individual transmission events and their timing is uncertain. The relationship between posterior times of infection and times of sampling is shown in Figure 5. There is approximately a 2-year gap between becoming infected and getting sampled.

Figure 5c shows how the estimated generation time varies over time. A large proportion of cases have generation time below 2 years, consistent with previous estimates in similar settings and in this outbreak [7, 2, 1, 15] The estimated mean generation time was lowest in and around 2004. However, not all posterior trees support the assumption that a given individual has infected someone else; in other words, there could be no transmission events between infection and sampling. Each point in Figure 5c is colored by the probability of the corresponding host ever infecting others. We see that many cases infected recently have a low probability of having infected others, and even if they do, the generation times tend to be short. This is in part due to censoring, as the sampling period ended. On the other hand, there are quite a few early cases that are very likely to be transmitters, and their generation times can be long (over 3 years). We note that due to the selection of closely related isolates for inclusion in the study, individuals who reactivated with TB strains that are not considered part of the outbreak are not shown here, and neither are infected individuals who did not become symptomatic before sampling ended. The times to sampling and to onward infection events reflect the censoring and case selection processes.

**Figure 5:**
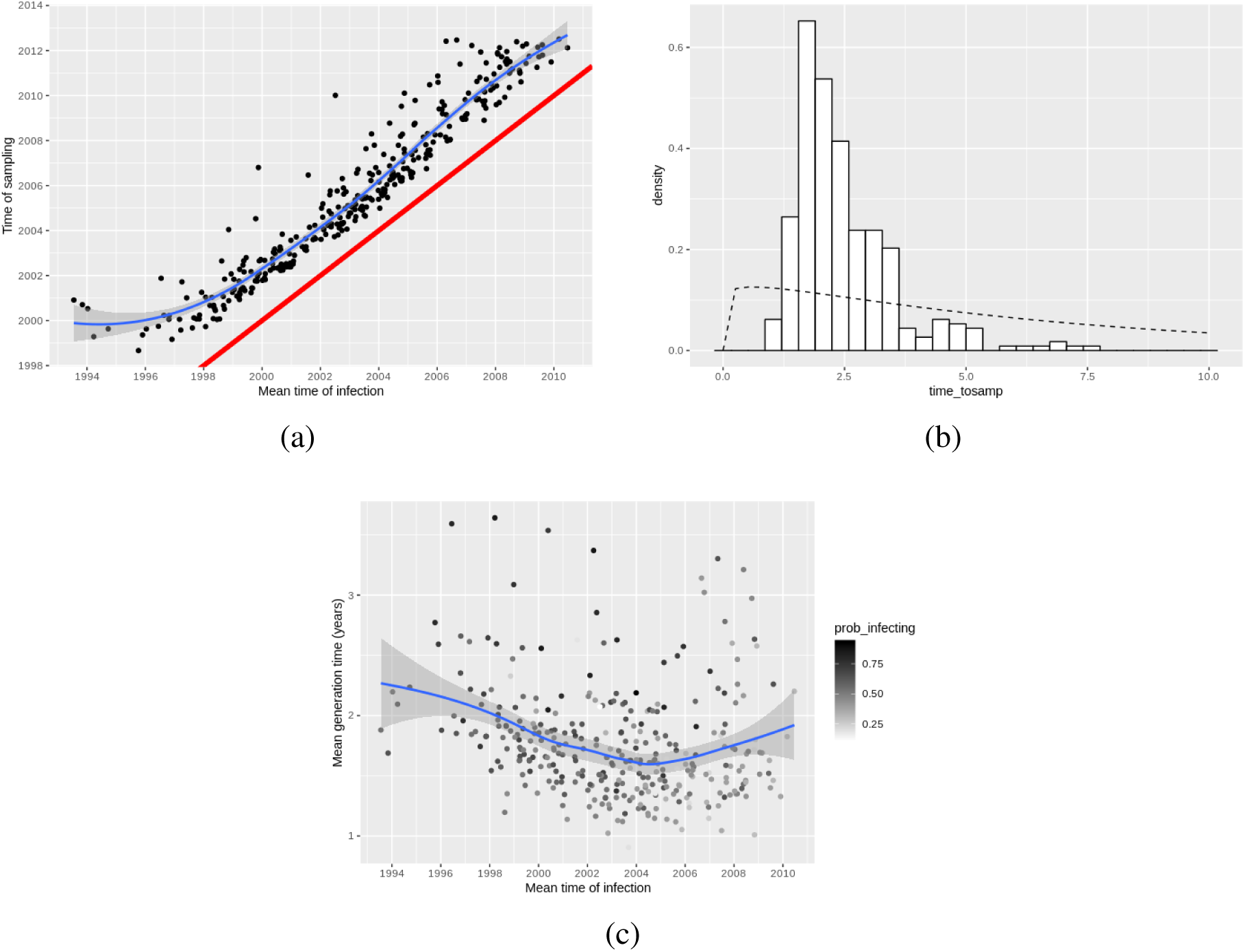
(a) Scatter plot of times of infection and sampling. Each dot corresponds to an individual host. A smooth line (blue) was fitted, a nd a r eference l ine (red) o f *y* = *x* w as a dded t o aid inspection. (b) Interval in years between times of infection and sampling in Figure 5, overlaid with the prior Gamma density used in TransPhylo (shape 1.1 and scale 6 years; dashed line). The few cases in the upper tail of the histogram correspond to cases earlier in the outbreak when sampling was poor. (c) Scatter plot of time of infection and generation time in years, each dot corresponds to an individual host. The individual cases are colored by their probability of infecting others, with darker color indicating a bigger probability. In (a) and (c), a smoother was fitted in order to better see the relation between the variables, using local polynomial regression fitting, or “loess”; the shaded area indicates 0.95 confidence interval level.

We sought to relate the covariate data to two outcomes from the outbreak reconstruction: whether an individual is likely to have transmitted TB, and whether an individual has a relatively long or short generation time (time between becoming infected and infecting others). Table 1 shows the aggregated performance of the unoptimised random forest classifiers in a confusion matrix. In the first task (identifying likely transmitters), we obtain an accuracy of 0.7, precision 0.72 and recall 0.83. A high recall rate is often desirable because it means that the probability of detecting cases that have infected others will be high. However in this case we have a false positive rate of 0.51, which means that one in two predicted positives will likely be a false alarm. This could still be helpful to case finding, as overall TB prevalence is low. Figure S3a shows the ROC curve for the first task, with area under curve (AUC) of 0.66.

**Table 1:**
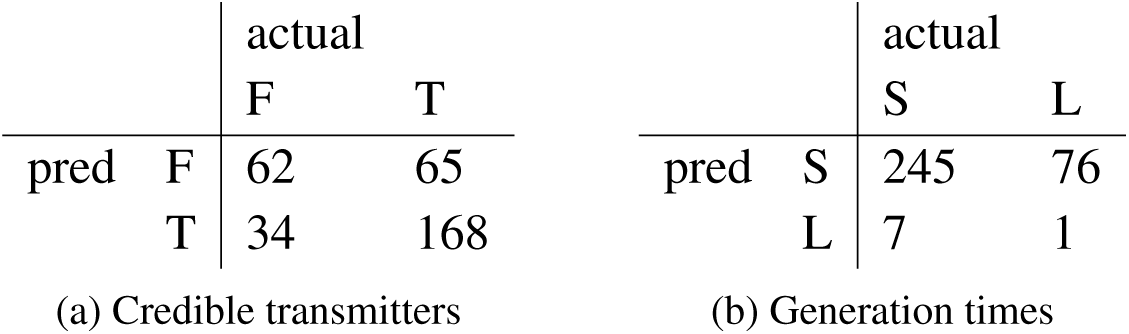
Confusion matrix of the random forest classifier on the validation set. Left: Classification: credible transmitter status. Right: Classification: long and short generation times.

Figure 6a shows feature importance from the random forest classifier. The importance of a variable is measured by the mean decrease of node impurity, in this case the Gini index, from splitting on that variable in the decision tree. If splitting on variable *A* reduces misclassification more than splitting on variable *B*, then *A* is considered to be more important than *B*. We find that the age of the patient is the most important variable by this measure, outweighing the other variables by a substantial margin. The importance findings may be interesting to epidemiologists offering insights into variables that may affect the likelihood of transmission. However, we should not be too confident because the classifier’s performance is not very good (although it is better than random guessing). Including additional covariates may improve classification. Our machine learning results also suggest that sputum smear status is not particularly important in predicting whether an individual transmitted TB in this outbreak.

**Figure 6:**
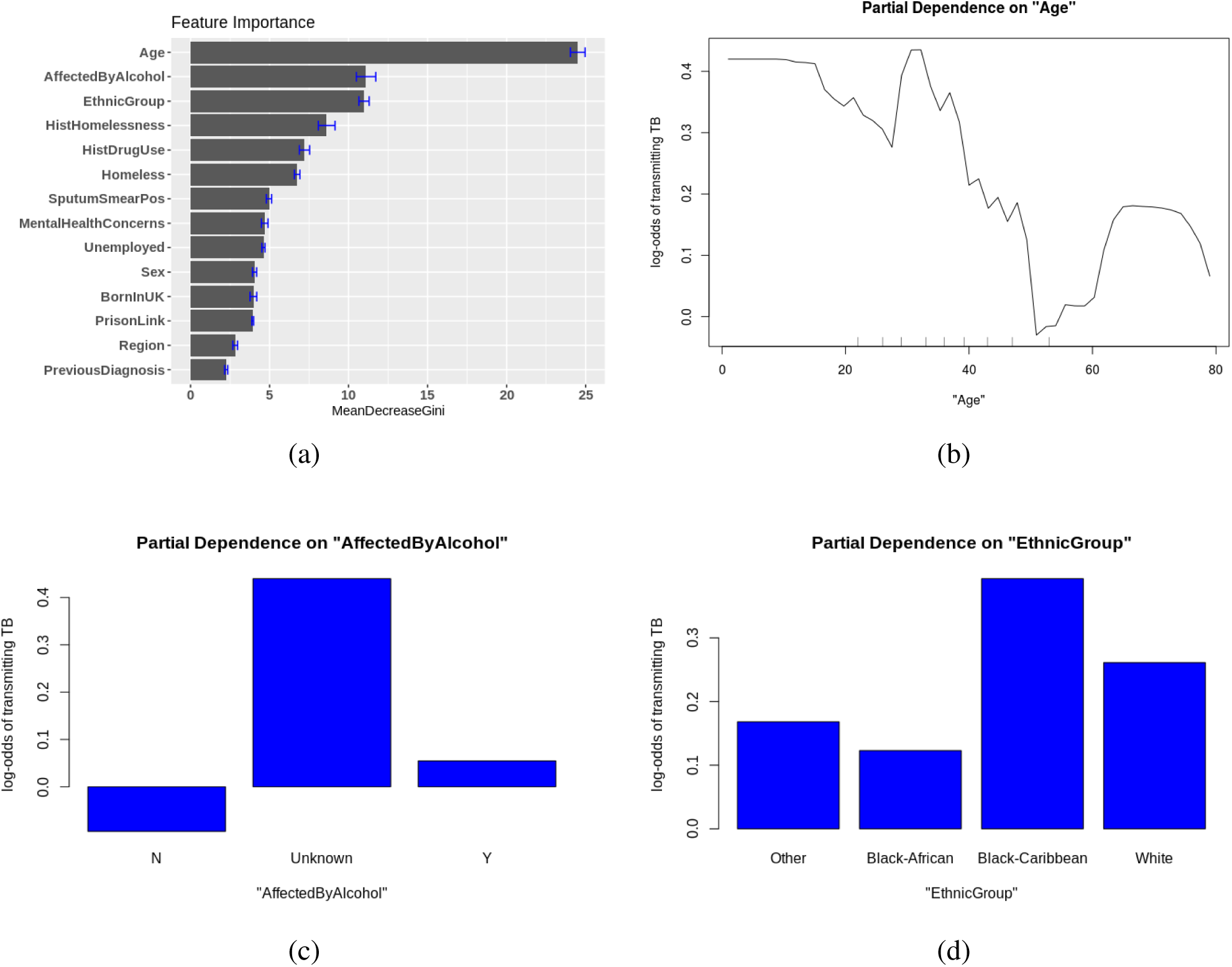
(a): Feature importance plot of the random forest model for classifying whether a host has transmitted TB to others. Importance is measured by the mean decrease in Gini index from splitting on the variable. The error bar is the standard error of the importance measure on 5 imputed datasets. (b)-(d): Partial dependence plot for age, alcoholism and ethnic group.

Partial dependence plots can be used to visualize the marginal effect of some predictors on the response variable, by averaging out the effects of all other variables. In Figure 6b-6d, we show the partial dependence on age, alcoholism and ethnic group, the three most important predictors, as log-odds. There is a sharp decrease in the likelihood of transmitting TB (according to the fitted model) after age 40; the odds increase somewhat in people over 60. We also note that the log-odds of transmitting TB is a little higher for those who are affected by alcohol than those who are not. The log-odds is largest if the alcoholism variable’s value is unknown, suggesting that there are more positive (Y; i.e. affected by alcohol) than negative (N) cases among those patients who had not reported their alcoholism history. We also observe that individuals of Black Caribbean heritage are more likely to appear as credible TB transmitters than other ethnic groups in our inference.

We also attempted to predict from the metadata whether an individual would have a long or short generation time. Treating the case where the generation time is more than 2 years as the ‘positive’ category, this classifier does not perform as well as trying to predict whether the host has transmitted TB, using the same set of covariates. The confusion matrix of this classifier is shown in Table 1b.

## 4. Discussion

We have demonstrated how transmission reconstruction from WGS data can be approached using Bayesian statistical inference of transmission trees, with data from a large TB outbreak in London. We have used a modified version of the TransPhylo approach to simultaneously infer transmission events on multiple trees, sharing parameters between them. This allows us to incorporate tree uncertainty into the transmission inference. The ways in which individuals live, work and interact is one of the driving forces for TB transmission [12]. Understanding the relationship between the covariates and transmission gives us insights into factors driving of TB transmission, and could provide guidance on effective control mechanisms to public health authority.

Although WGS can be insufficient to resolve transmission chains due to lack of detectable variation between cases [5, 4], our statistical approach can refine the analysis usually used in outbreak investigations. We identified more transmission events with reasonable confidence than those that were suggested directly by the data [5]. When patient-level covariates such as demographic and clinical data are available, machine learning algorithms can be used to predict individual-level variables (ie “credible transmitter” status) derived from transmission reconstruction, providing a means to assess the importance of the covariates for these quantities.

With the move to routine WGS of all TB isolates by Public Health England, it is important to understand the role WGS data can play in outbreak investigations and in understanding transmission. With current sequencing technology and variant-calling pipelines, WGS data may contain insufficient variation to reconstruct individual transmission events with high confidence. It may be that variation simply does not occur rapidly enough in TB to obtain much more information about direct transmission, making the development of approaches to better integrate additional epidemiological data very important [4]. However, sequence data can still contribute to epidemiological analysis through the kind of integrative analysis we have done here as well as through refuting putative direct transmission events when the relevant isolates are very distinct genomically. It is possible that new longer-read technologies and improved variant calling may ultimately allow us to capture additional variation occurring in repeat regions, hyper-variable regions, or variation due to insertions and deletions; this would likely be helpful in epidemiological investigations of TB outbreaks in a range of settings.

Our approach has some significant limitations. It is a three-stage approach: construction of timed phylogenetic trees, transmission analysis, followed by machine learning to connect the demographic and clinical data to the transmission analysis. While we have made efforts to take uncertainty into account at each stage by, for example, simultaneously analysing 50 posterior timed phylogenetic trees, joint estimation of the transmission trees and phylogenetic trees together might be preferable if it could be done in a practical way. There would also be advantages to developing statistical and modelling tools to directly (and simultaneously with the phylogeny and transmission trees) estimate the contributions of each covariate to transmissibility, speed of progression of disease and other factors. Instead, here we assumed a ‘ground truth’ which was in fact estimated with TransPhylo. Developing the appropriate inference tools would require overcoming the challenge of handling unsampled cases (and the unknown cases they may have infected, and so on) despite the unknown covariate data for the unknown cases. Currently, the mathematics at the heart of TransPhylo does not naturally allow for a likelihood model that extends in this way.

It would additionally help to analyse sequence data together with outbreak control efforts in real time [18, 10]. In tuberculosis, with outbreaks lasting years, this is very feasible. Results could inform the outbreak investigation by directing attention towards individuals without a probable infector (and thus a likely contact of an unknown case), by informing public health bodies as to how quickly cases need to be found to interrupt transmission and towards communities or sub-groups with higher numbers of estimated unsampled cases nearby in the transmission tree. To take these actions would require relatively rapid WGS and analyses, but this is now increasingly feasible [9]. WGS data can readily be used to refute transmission events, and routine sequencing has the potential to lead to dramatic improvements in understanding and treating resistant disease, particularly if genome-based resistance predictions can be made quickly enough to inform treatment [9].

One recurring message [5, 7, 4] is that WGS data alone are likely to be insufficient for reconstructing individual transmission events; however, our statistical approach can improve the analysis of WGS data together with covariates, and uncover patterns of transmission. Multiple data sources are required to obtain the best possible understanding of transmission events and transmission patterns. At least with current sequencing and bioinformatics pipelines, clinical, contact, epidemiological and demographic data cannot be replaced with sequencing even though WGS data can have a significant role to play.

## 5. Supplementary Materials

Figure S1 shows the trajectories of model parameters during the TransPhylo MCMC run. Figure S2 shows the estimated number of unsampled cases, and Figure S3 shows the receiver-operator characteristic curves for the classification tasks. A random classifier would have an area under the curve (AUC) on these plots of 0.5. Our results suggest that the covariates are better predictors of whether an individual transmits TB than of whether an individual has a longer-than-usual generation time, but we note that these results use the TransPhylo estimates in place of the true status (infector; generation time), as the truth is not known.

**Figure S1:**
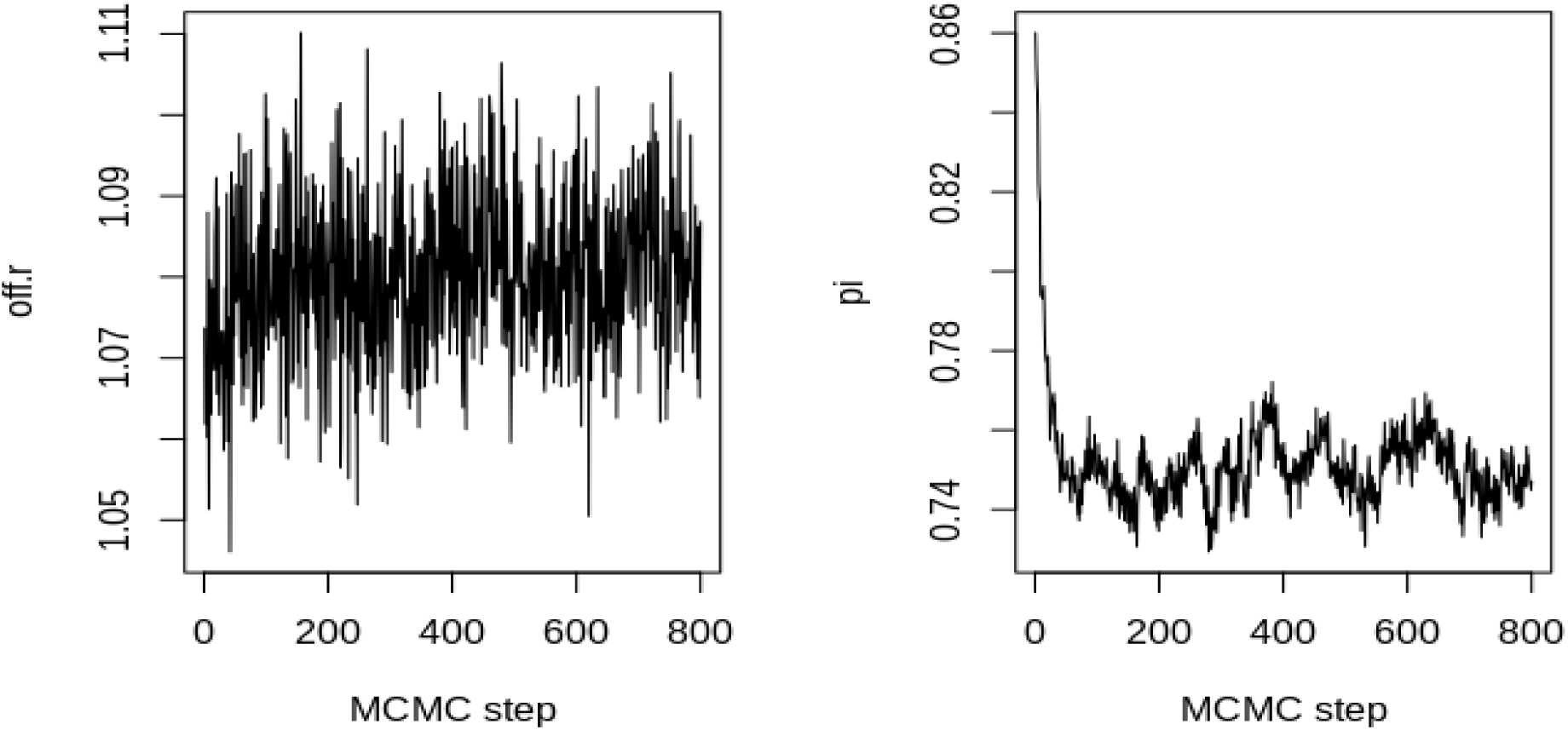
Trace plot of model parameters. neg — within-host diversity, off.r — first parameter of negative binomial offspring distribution, or equivalently the basic reproduction number, and pi — sampling rate.

**Figure S2:**
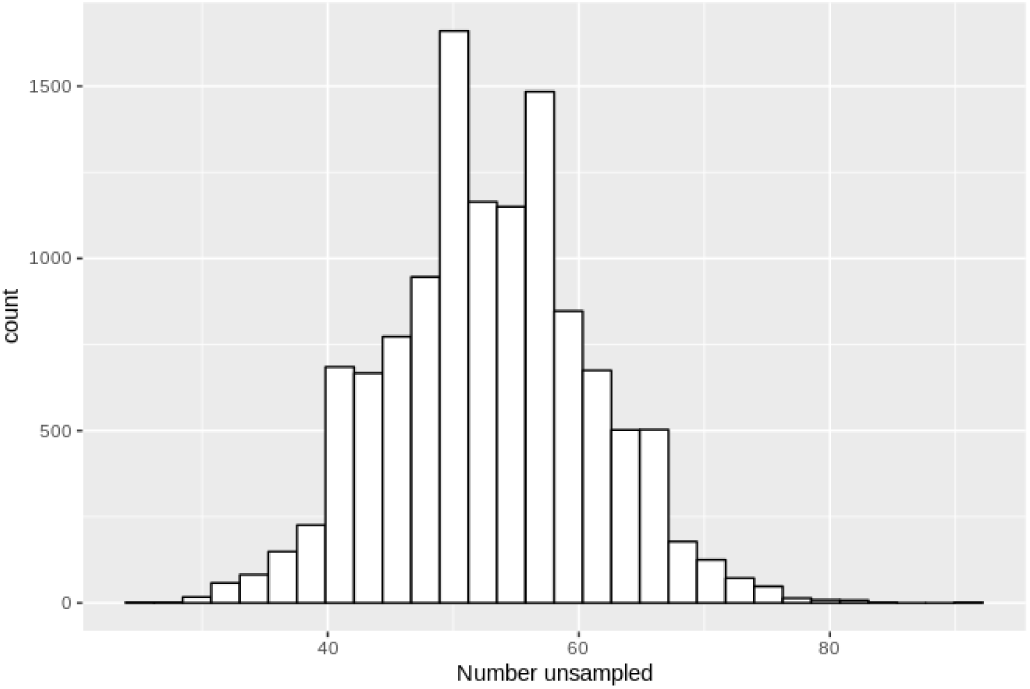
Histogram of number of unsampled cases, generated from the combined posterior transmission trees.

**Figure S3:**
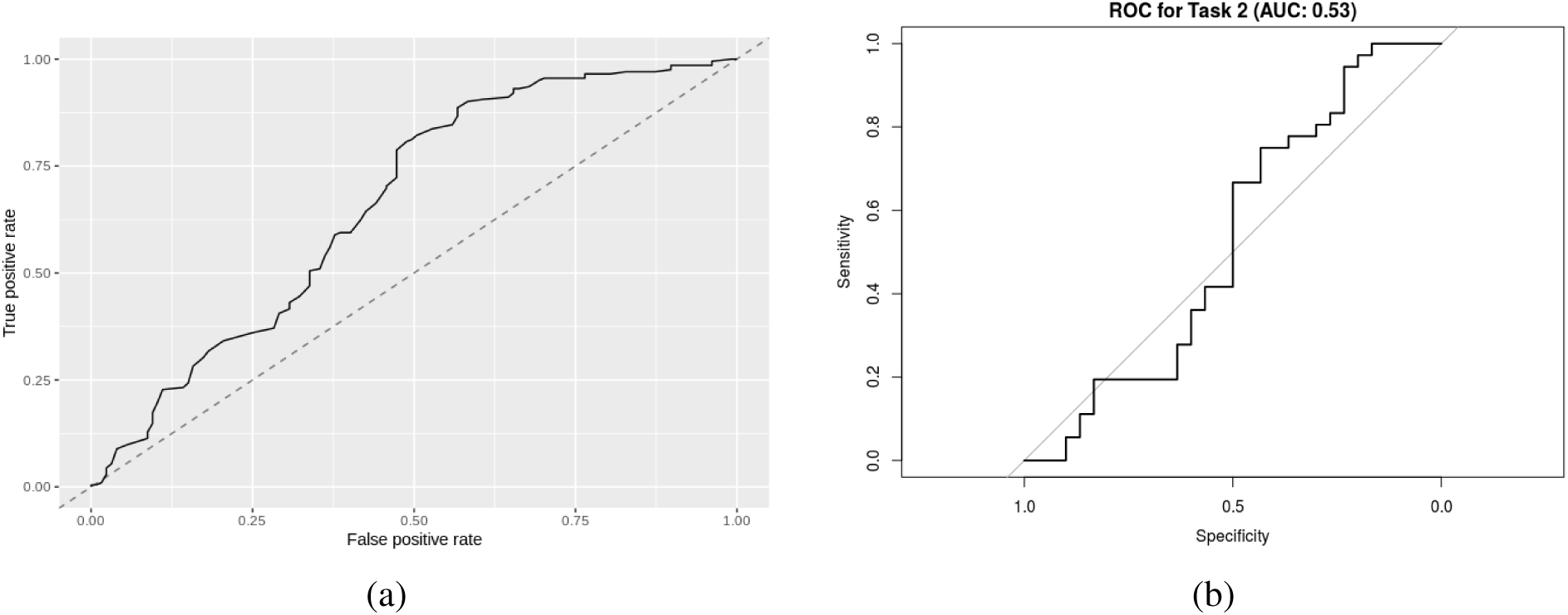
ROC curves illustrating the overall performance of the classifiers. A random classifier is expected to have an AUC (area under the curve) of 0.5 and perfect classification, with no false positives or false negatives, achieves an AUC of 1. (a): Task one: predicting if a host has transmitted TB. (b) Task two: predicting if a host has over 2 years generation time.

**Figure S4:**
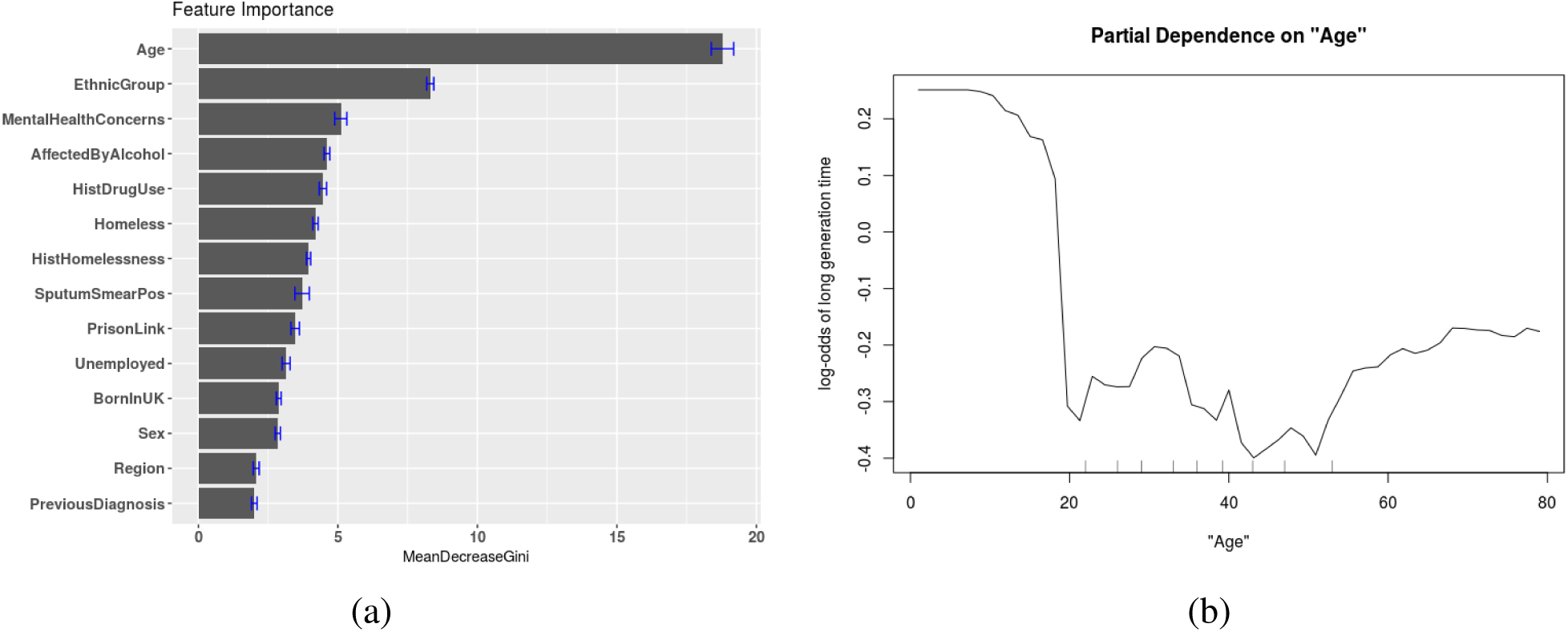
(a): Feature importance plot of the random forest model for classifying whether a host has generation time greater than 2 years. Importance is measured by the mean decrease in Gini index from splitting on the variable. The error bar is the standard error of the importance measure on 5 imputed datasets. (b): Partial dependence plot for age variable.

## References

[1] Diepreye Ayabina, Janne O Ronning, Kristian Alfsnes, Nadia Debech, Ola B Brynildsrud, Trude Arnesen, Gunnstein Norheim, Anne-Torunn Mengshoel, Rikard Rykkvin, Ulf R Dahle, Caroline Colijn, and Vegard Eldholm. Genome-based transmission modelling separates imported tuberculosis from recent transmission within an immigrant population. Microbial Genomics, September 2018.

[2] Marcel A Behr, Paul H Edelstein, and Lalita Ramakrishnan. Revisiting the timetable of tuberculosis. BMJ, 362, 2018.

[3] Remco Bouckaert, Joseph Heled, Denise Kühnert, Tim Vaughan, Chieh-Hsi Wu, Dong Xie, Marc A. Suchard, Andrew Rambaut, and Alexei J. Drummond. Beast 2: A software platform for bayesian evolutionary analysis. PLOS Computational Biology, 10(4):1–6, 04 2014.

[4] Finlay Campbell, Camilla Strang, Neil Ferguson, Anne Cori, and Thibaut Jombart. When are pathogen genome sequences informative of transmission events? PLoS Pathogens, 14(2):e1006885, February 2018.

[5] Nicola Casali, Agnieszka Broda, Simon R Harris, Julian Parkhill, Timothy Brown, and Francis Drobniewski. Whole genome sequence analysis of a large isoniazid-resistant tuberculosis outbreak in london: a retrospective observational study. PLoS medicine, 13(10):1–18, 2016.

[6] Caroline Colijn and Jennifer Gardy. Phylogenetic tree shapes resolve disease transmission patterns. Evolution, Medicine, and Public Health, 2014(1):96–108, 2014.

[7] Xavier Didelot, Christophe Fraser, Jennifer Gardy, and Caroline Colijn. Genomic infectious disease epidemiology in partially sampled and ongoing outbreaks. Molecular biology and evolution, 2017.

[8] Xavier Didelot, Jennifer Gardy, and Caroline Colijn. Bayesian Inference of Infectious Disease Transmission from Whole-Genome Sequence Data. Molecular Biology and Evolution, 31(7):1869–1879, 04 2014.

[9] Ronan M Doyle, Carrie Burgess, Rachel Williams, Rebecca Gorton, Helen Booth, James Brown, Josephine M Bryant, Jackie Chan, Dean Creer, Jolyon Holdstock, Heinke Kunst, Stefan Lozewicz, Gareth Platt, Erika Yara Romero, Graham Speight, Simon Tiberi, Ibrahim Abubakar, Marc Lipman, Timothy D McHugh, and Judith Breuer. Direct Whole-Genome Sequencing of Sputum Accurately Identifies Drug-Resistant Mycobacterium tuberculosis Faster than MGIT Culture Sequencing. J. Clin. Microbiol., 56(8), August 2018.

[10] Jennifer L Gardy and Nicholas J Loman. Towards a genomics-informed, real-time, global pathogen surveillance system. Nat. Rev. Genet., 19(1):9–20, January 2018.

[11] H Maguire, S Brailsford, J Carless, M Yates, L Altass, S Yates, S Anaraki, A Charlett, S Lozewicz, M Lipman, and G Bothamley. Large outbreak of isoniazid-monoresistant tuberculosis in London, 1995 to 2006: case-control study and recommendations. Euro Surveill., 16(13), March 2011.

[12] Barun Mathema, Jason R Andrews, Ted Cohen, Martien W Borgdorff, Marcel Behr, Judith R Glynn, Roxana Rustomjee, Benjamin J Silk, and Robin Wood. Drivers of tuberculosis transmission. The Journal of infectious diseases, 216:S644–S653, 2017.

[13] F Neely, H Maguire, F Le Brun, A Davies, D Gelb, and S Yates. High rate of transmission among contacts in large London outbreak of isoniazid mono-resistant tuberculosis. J. Public Health, 32(1):44–51, March 2010.

[14] World Health Organization. Standards and benchmarks for tuberculosis surveillance and vital registration systems: checklist and user guide. Technical report, World Health Organization, 2014. WHO/HTM/TB/2014.2 and WHO/HTM/TB/2014.6 (Checklist).

[15] Andreas Roetzer, Roland Diel, Thomas A. Kohl, Christian Rückert, Ulrich Nübel, Jochen Blom, Thierry Wirth, Sebastian Jaenicke, Sieglinde Schuback, Sabine Rüsch-Gerdes, Philip Supply, Jörn Kalinowski, and Stefan Niemann. Whole genome sequencing versus traditional genotyping for investigation of a mycobacterium tuberculosis outbreak: A longitudinal molecular epidemiological study. PLOS Medicine, 10(2):1–12, 02 2013.

[16] M C Ruddy, A P Davies, M D Yates, S Yates, S Balasegaram, Y Drabu, B Patel, S Lozewicz, S Sen, M Bahl, E James, M Lipman, G Duckworth, J M Watson, M Piper, F A Drobniewski, and H Maguire. Outbreak of isoniazid resistant tuberculosis in north London. Thorax, 59(4):279–285, April 2004.

[17] G Satta, M Lipman, G P Smith, C Arnold, O M Kon, and T D McHugh. Mycobacterium tuberculosis and whole-genome sequencing: how close are we to unleashing its full potential? Clin. Microbiol. Infect., 24(6):604–609, June 2018.

[18] Patrick Tang and Jennifer L Gardy. Stopping outbreaks with real-time genomic epidemiology. Genome Medicine, 6(11):104, November 2014.

[19] Grant Theron, Helen E Jenkins, Frank Cobelens, Ibrahim Abubakar, Aamir J Khan, Ted Cohen, and David W Dowdy. Data for action: collection and use of local data to end tuberculosis. Lancet, October 2015.

[20] Chongguang Yang, Tao Luo, Xin Shen, Jie Wu, Mingyu Gan, Peng Xu, Zheyuan Wu, Senlin Lin, Jiyun Tian, Qingyun Liu, ZhengAn Yuan, Jian Mei, Kathryn DeRiemer, and Qian Gao. Transmission of multidrug-resistant mycobacterium tuberculosis in shanghai, china: a retrospective observational study using whole-genome sequencing and epidemiological investigation. The Lancet Infectious Diseases, 17(3):275 – 284, 2017.

